# Attentional rhythmic blink: Theta/Alpha balance in neural oscillations determines the rhythmicity in visual sampling

**DOI:** 10.1101/2022.04.15.488436

**Authors:** Tomoya Kawashima, Masamichi J Hayashi, Kaoru Amano

## Abstract

Brain oscillations in the theta (3–7 Hz) and alpha (7–13 Hz) bands are implicated in visual perception and attention. We show that in an attentional blink paradigm, where the task requires detecting two targets presented in rapid succession, perceptual performance varied with the rhythms at these two frequencies, which we name attentional rhythmic blink. In the absence of distractors, second target detection performance fluctuated at the theta rhythm, but the fluctuation frequency shifted toward alpha rhythm when distractors were interspersed with the targets. We further show, in magnetoencephalography experiments, that a change in the dominant frequency of ongoing neural oscillations accompanied those in perceptual performance, with the parietal theta being more pronounced in the no-distractor and the occipital alpha in the distractor conditions, respectively. We propose that perceptual rhythms may depend on the power balance between ongoing neural oscillations, determined by the task-specific demand.

## Introduction

Our sensory experience feels seamless. However, recent studies have suggested that our brain samples sensory information rather rhythmically over time, which has been understood to reflect neural oscillations (Buzsáki, 2006; Fiebelkorn & Kastner, 2019, 2020; Fries, 2015; VanRullen, 2016). Although both alpha (7–13 Hz) and theta (4–7 Hz) oscillations seem to be crucial for visual perception and attention, we do not yet understand how the frequency in rhythmic sampling is determined. Empirical studies have shown that ongoing occipital alpha oscillations determine fluctuations in visual sensitivity (Busch et al., 2009; Busch & VanRullen, 2010; Mathewson et al., 2009; VanRullen, Carlson, & Cavanagh, 2007). Alpha oscillations are also known to be related to visual attention: directing attention to the left or right hemifield suppresses contralateral and enhances ipsilateral alpha oscillations (Kelly et al., 2006; Jensen & Mazaheri, 2010; Worden et al., 2000; Thut et al., 2006). In addition, involvement of theta oscillations has been repeatedly observed in spatial attention (Davidson et al., 2018; Dugué et al., 2017; Fiebelkorn et al., 2013: Holcombe & Chen, 2013; Landau & Fries, 2012; Michel et al., 2021; Senoussi et al., 2019; Song et al., 2014) and feature-based attention tasks (Re et al., 2019).

To investigate the role of oscillations in visual perception, we used the attentional blink paradigm in a visual detection task (Raymond et al., 1992). Subjects needed to detect two successively presented targets. However, their ability to detect the second target was variably impaired, depending on the elapsed time between the presentation of the two targets. In these psychophysical experiments, we demonstrated a rhythmicity of behavioral performance driven by attention to temporally distributed items, which we name *attentional rhythmic blink*. Importantly, the fluctuation frequency of the second target detection performance changed depending on whether task-irrelevant items (distractors) were temporally interspersed with the targets; while performance fluctuated at the theta rhythm when distractors were absent, the fluctuation shifted toward the faster frequencies of the alpha rhythm when distractors were presented. The enhancement of alpha oscillations has been shown to have a critical role for suppressing task-irrelevant visual processing (Clayton et al., 2018; Jensen & Mazaheri, 2010; Klimesch et al., 2007; Mathewson et al., 2011). Based on these observations, we hypothesized that the two perceptual rhythms may originate from the change in the balance of neural oscillations. To test this hypothesis, we measured the neural correlations of these behavioral oscillations using magnetoencephalography (MEG). We found that, just before the appearance of the second target, the power of parietal theta was dominant over the power of occipital alpha oscillations in the no-distractor conditions. The converse, occipital alpha dominant over parietal theta, was true in the distractor condition. Moreover, the phases of the neural oscillations were predictive of whether the observers could detect the second target.

Taken together, our results suggest that perceptual rhythm is not constant but, rather, dynamically modulated by the power balance between theta and alpha oscillations affected by task demands, such as distractor suppression. This explanation of frequency change in rhythmic sampling may provide a unified framework for attentional rhythms in spatial, feature-based, and temporal attention in vision.

## Results

### Attentional rhythmic blink at the theta frequency

The involvement of theta rhythm in behavioral performances has been reported not only in spatial attention tasks (e.g., Fiebelkorn et al., 2013) but also in feature-based attention tasks (Re et al., 2019), which might reflect general oscillatory mechanisms in multi-item attention (Huang et al., 2015). We investigated whether such rhythmicity is implicated in the temporal domain. Temporal attention is required for selecting a target embedded in a temporal sequence of multiple items that are presented at the same location in rapid succession (e.g., Jiang & Chun, 2001). Utilizing a paradigm called attentional blink, which allows assessing the temporal aspects of visual attention by removing the effect of spatial and feature-based attention, we unveiled rhythmic variations of attentional blink at both theta and alpha frequencies, which we name *attentional rhythmic blink*.

In attentional blink, where participants detect two targets, T1 and T2, briefly separated in time (Fig. 1a), detection of the second target (T2) is deteriorated for T1-to-T2 lag of approximately 300 ms (Chun & Potter, 1995; Dux & Marois, 2009; Raymond et al., 1992). To investigate the dependence of the behavioral performance in the attentional blink task on the duration of the T1-to-T2 lag, we randomly sampled the lag in discrete 20 ms increments (corresponding to 50 Hz sampling rate) on repeated trials and measured the performance in 31 subjects (Experiment 1a). To assess the temporal profile of attentional blink performances, we calculated the T2 conditional accuracy given that T1 was correct (T2|T1-correct) as a function of the temporal lag between T1 and T2. As shown in Figs. 2a and 2b, there was a trend that T2 detection accuracy was worse for T1-to-T2 lag of 300 ms (67% correct) than for a lag of 800 ms (71% correct) (*t* (30) = 2.00, *p* = 0.054, *d* = 0.36), replicating the conventional attentional blink phenomenon. More interestingly, the T2 detection accuracy showed a cyclic fluctuation as a function of T1-to-T2 lag (Fig 2a and 2c). To measure this periodicity in T2 detection, we performed the Fast Fourier transform (FFT) analysis to convert the T2 accuracy function in the temporal (T1-to-T2 lag) domain into the temporal frequency domain (see Methods). Spectrum characteristics of the T2 detection accuracy (Fig. 2d) indicated behavioral oscillations at around 4 Hz (3.6–4.7 Hz, *p* < 0.05, corrected for multiple comparison). This result suggests that behavioral performance fluctuated while attention was temporally allocated to multiple items, which is in line with recent findings utilizing spatial-based or feature-based attention paradigm (e.g., Huang et al., 2015; Fiebelkorn et al., 2013: Re et al., 2019). A similar theta-fluctuation of T2-detection was observed for T2|T1-error trials (Fig. S1), suggesting that the theta sampling of two temporally separated items does not depend on whether the first item was correctly encoded in working memory.

**Fig. 1:**
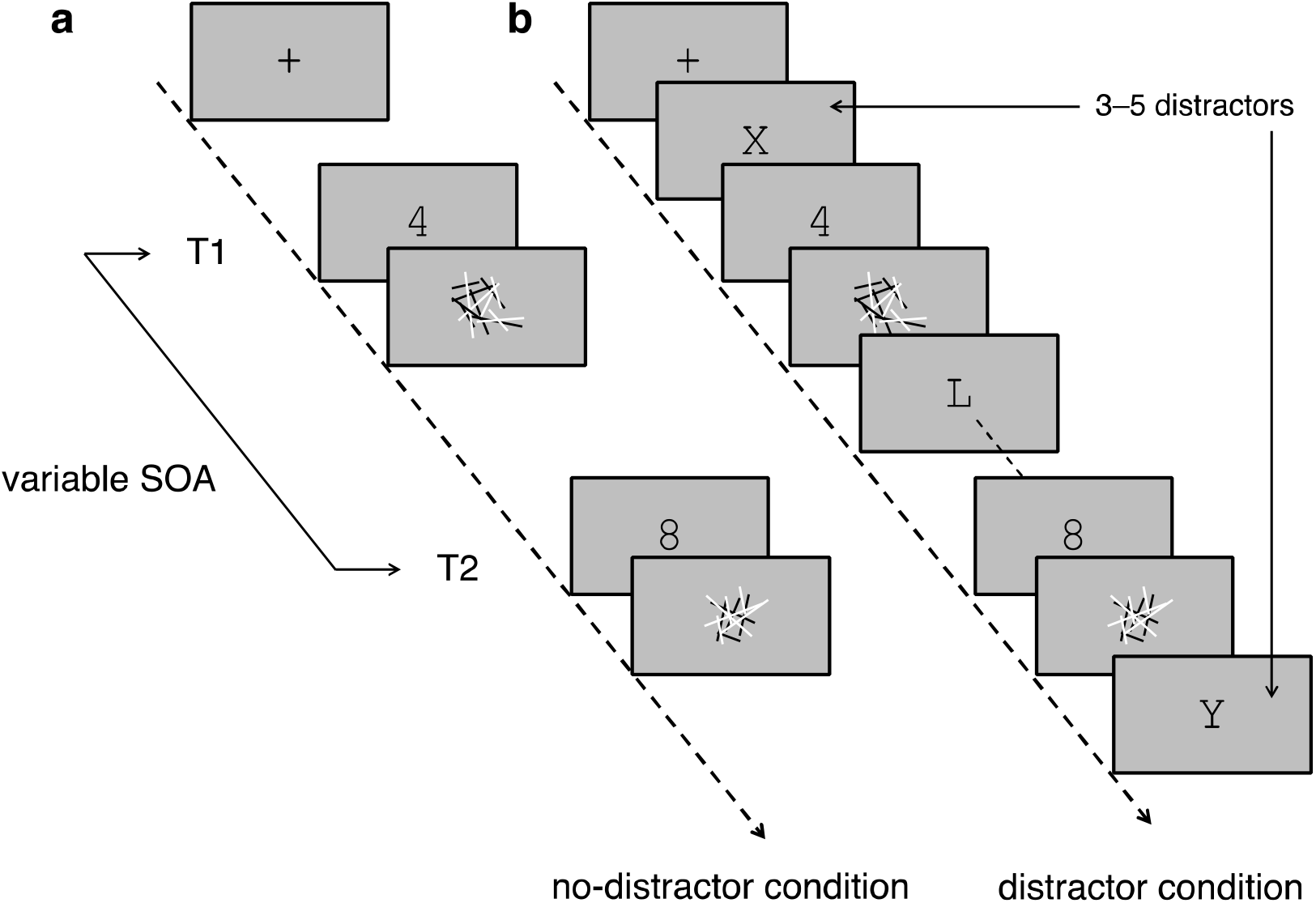
Schematics of experimental trials. In both behavioral and MEG experiments, the task was to detect two targets (single digits) successively presented with a variable stimulus onset asynchrony (SOA) sampled in the 100 to 1000 ms range. Both targets were masked with black and white line-fragments. **a** No-distractor condition, where each trial only included the two targets. **b** Distractor condition, where in each trial the two targets were embedded in a rapid serial visual presentation (RSVP) stream that included a variable number of distractors (single letters).

**Fig. 2:**
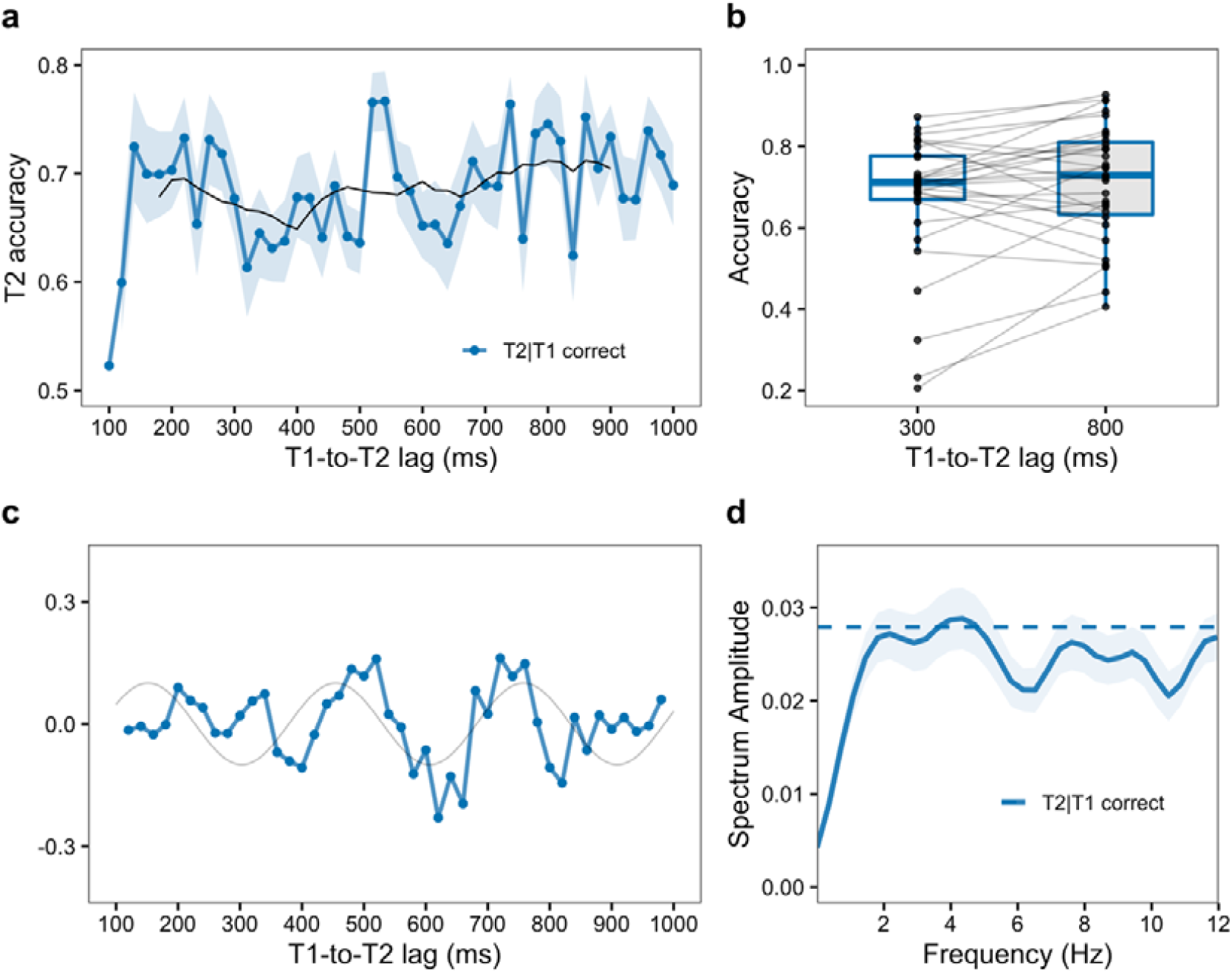
Attentional rhythmic blink in the no-distractor condition (Experiment 1a: *n* = 31). **a** T2 detection accuracy (mean ± SEM) in T2|T1-correct trials, as a function of T1-to-T2 lag (100 ms to 1000 ms interval discretized 20-ms steps), averaged across participants (blue line). Smoothing, using 10-point moving average, was applied to determine the classic attentional blink phenomena (black line). **b** The mean T2 detection accuracy for T1-to-T2 lags of 300 ms and 800 ms, obtained after smoothing separately for each trial type. Each dot represents the mean accuracy for a single participant. **c** Detrended accuracy time course for one participant. Note the oscillatory pattern over time. 3-point moving average was applied for visualization. The gray curve shows the sine curve at the peak frequency (3.3 Hz; amplitude and phase are arbitrary) in the Fourier spectrum for this subject. **d** Grand average spectrum (mean ± SEM) for detrended accuracy time courses as a function of frequency for the T2|T1-correct condition. A dashed line indicates the threshold statistical significance (*p* < 0.05) after permutation test and corrections for multiple comparisons.

### Attentional rhythmic blink at the alpha frequency

Next, we assessed whether the attentional rhythmic blink depended on the existence of distractors interspersed between two targets (Experiment 1b). Participants (*n* = 28) were asked to select two target numbers while ignoring task-irrelevant letters (Fig. 1b). The results (Fig. 3a & b) again replicated conventional effects of attentional blink: 31% correct at 300 ms vs. 43% correct at 800 ms (*t* (27) = 6.56, *p* < 0.001, *d* = 1.24). Relatively small AB magnitude compared to the original task (e.g., Shapiro et al., 1997), would be due to the arrhythmic RSVP stream employed in the current study, which is known to reduce the AB magnitude (Martin et al., 2011). More importantly, as shown in Fig. 3d, detection accuracy fluctuated at approximately 8 Hz (7.2–9.1 Hz: *p* < 0.05, corrected for multiple comparison). These results indicate that behavioral performance cyclically fluctuates when participants must detect two temporally separated targets, and the fluctuation frequency depends on whether distractors are presented or not. In contrast alpha-band fluctuations observed in the T2|T1-correct trials, T2 accuracy in the T2|T1-error trials fluctuated at theta frequencies. This difference possibly suggests that successful encoding of the first item is crucial for the alpha sampling (Fig S2: see Discussion for detail).

**Fig. 3:**
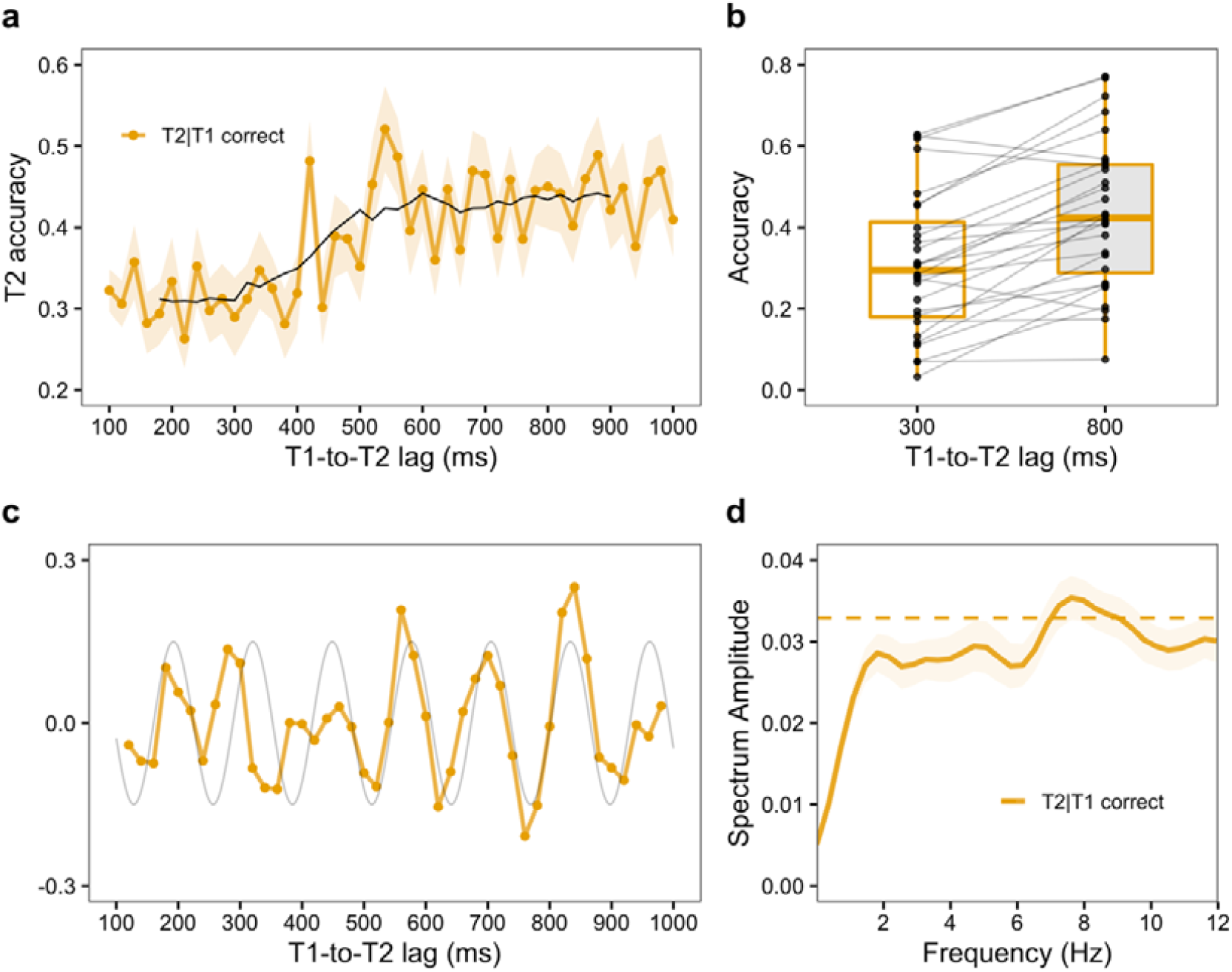
Attentional rhythmic blink in the distractor condition (Experiment 1b: *n* = 28). **a** T2 detection accuracy (mean ± SEM) as a function of T1-to-T2 lag for T2|T1-correct trials, averaged across participants (yellow line). Smoothing (10-point moving average) was applied to determine the classic attentional blink phenomenon (black line). **b** The mean T2 detection accuracy for T1-to-T2 lags of 300 ms and 800 ms, obtained after 10-point moving average for each trial type. At each lag, one dot represents the mean accuracy for each participant. **c** Detrended accuracy time course for one participant. Note the temporal fluctuation over lag-time. 3-point moving average was applied for visualization. The gray curve shows the sine curve at the peak frequency (7.8 Hz; amplitude and phase are arbitrary) in the Fourier spectrum for this subject. **d** Grand average spectrum (mean ± SEM) for detrended accuracy time courses as a function of frequency for the T2|T1-correct condition. The dashed line indicates the threshold statistical significance (*p* < 0.05) after permutation test and corrections for multiple comparisons.

### Neural correlates of attentional rhythmic blink

When participants needed to detect two temporally adjacent targets, the detection accuracy of the second target periodically fluctuated as a function of the interval between the targets. Furthermore, the frequency depended on the existence of distractors. Namely, the fluctuation frequency was shifted from ∼4 Hz in the no-distractor condition (Experiment 1a) to ∼8 Hz in the distractor condition (Experiment 1b). The results obtained in Experiment 1 for rhythmic blink suggest that sampling frequency in attention to temporally distributed items is sensitive to the existence of distractors. Based on the substantial evidence implicating the role of alpha oscillations for suppressing distractors (e.g., Jensen & Mazaheri, 2010), we hypothesize that distractors added in the visual streams might have altered the power balance between theta and alpha oscillations, which in turn resulted in the change in behavioral oscillations. To test this hypothesis, we measured MEG signals while participants performed an attentional blink task with and without distractors in Experiment 2 (*n* = 27). The corresponding behavioral data are shown in Fig.S3.

We first analyzed how the power of neural oscillations prior to the time of T2 presentation is modulated by the presence of distractor stimuli (Fig 4; see Fig S4 for both hemispheres). Since behavioral oscillations are thought to originate from ongoing alpha oscillations in the early visual cortex and theta oscillations in higher-order attentional areas (Dugué & VanRullen, 2017; Helfrich et al., 2017, 2018; Fiebelkorn et al., 2019), we focused our analysis on corresponding ROIs in the lateral occipital and superior parietal areas in the left and right hemispheres. We examined the differences, presence versus absence of distractors, in the theta and alpha oscillations preceding the time of T2 presentation. We found that the pre-stimulus theta (4-7 Hz) power was significantly higher for the no-distractor condition in the superior parietal region (Fig 4a, left). The difference was found only in the left hemisphere, not in the right hemisphere (Fig S4). This lateralization is consistent with several neuroimaging studies showing that allocating attention to temporally distributed items is associated with the left parietal cortex (Coull & Nobre, 1998; Griffin et al., 2001). In contrast, the pre-stimulus alpha (8–10 Hz) power was significantly higher for the distractor condition in the lateral occipital regions of both hemispheres (Fig 4a and Fig S4). These results suggest that the presence of distractors affected the relative power balance between theta and alpha oscillations: occipital alpha is dominant when distractors were required to be suppressed, while parietal theta is dominant when no distractors are presented. In addition to these changes of power in the theta and alpha bands, we also found an increase in the beta power prior to the T2 presentation in both ROIs, possibly reflecting the neural preparation for the upcoming stimuli (Meijer et al., 2016).

**Fig. 4:**
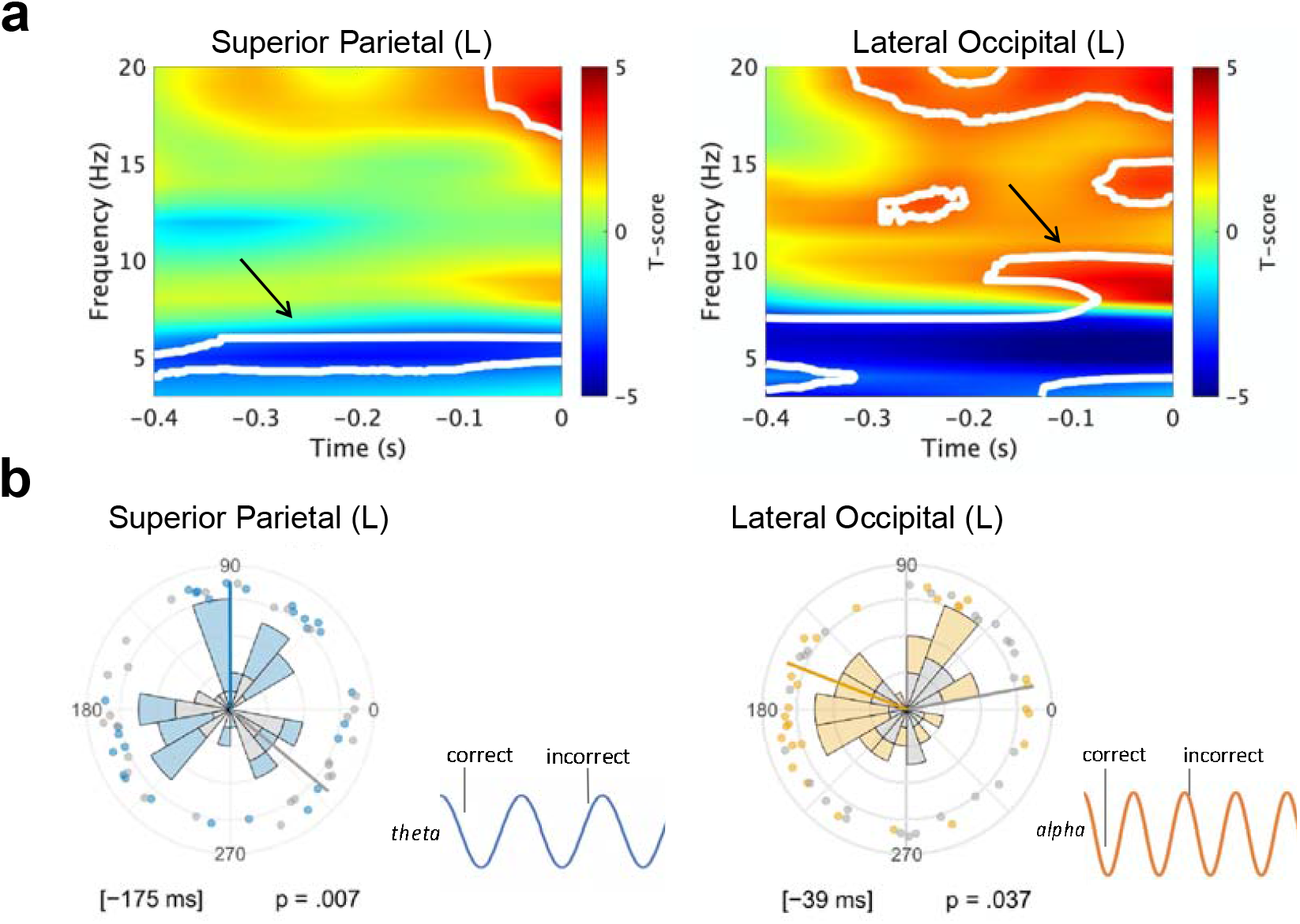
Pre-stimulus oscillatory power difference between distractor vs. no-distractor conditions and phase differences between correct vs. incorrect trials in Experiment 2. **a** The difference in the pre-stimulus power of neural oscillations between distractor vs. no-distractor conditions in left superior parietal (left) and left lateral occipital ROIs (right). The positive *t*-score values show that the power was larger for the distractor condition than for the no-distractor condition. White contours denote significant cluster-corrected effects (*p* < 0.05, FDR corrected). The arrows indicate pre-stimulus changes in the power in theta (4–7 Hz) and alpha (8–10 Hz) bands due to the presence of distractors. **b** The phase of oscillations in the theta-band (5–6 Hz: left) and alpha-band (8–10 Hz: right) prior to T2 presentation in the no-distractor (left panel) and distractor conditions (right panel). Data from correct trials shown in color; those in incorrect trials, in gray. The phases were referenced to the time point of maximal significance of the power (−175 ms for theta; and −39 ms for alpha). Phase differences were tested using a circular Watson-Williams test (thick radial lines represent the mean phase angle across participants). Anti-phase relationships between correct and error trials were observed in both the parietal theta and lateral occipital alpha oscillations. Each dot represents the mean angle for one participant.

The phase of pre-stimulus theta or alpha oscillations have been reported to affect visual perception (e.g., Han & VanRullen, 2017; van Dijk et al., 2008). This motivated us to compare the phase distributions between correct and incorrect trials using Watson-Williams circular test (Berens, 2009). The reference time point for phase determination was based on the minimum *p*-values obtained in the pre-T2 power analysis (Dugué et al., 2011). This analysis showed that, for theta oscillations (5–6 Hz; Figure 4b, left) measured in the left superior parietal region, significant phase difference between correct and incorrect trials was observed only in the no-distractor condition (130°, *F* (1, 53) = 8.01, *p* = 0.007). In contrast, for the alpha-band (8–10 Hz; Figure 4b, right), significant phase difference was observed in the left occipital area only in the distractor condition (148°, *F* (1, 53) = 4.57, *p* = 0.037). Fig. S2 shows a similar observation made in the right occipital area (195°, *F* (1, 53) = 7.91, *p* = 0.007). These results suggest that the phase of pre-stimulus parietal theta and occipital alpha oscillations, whose power was modulated by distracting items, affect the attentional rhythmic blink.

## Discussion

Recent studies have shown that our visual system samples information rhythmically. Investigation of the factors determining the frequency in this perceptual rhythm is crucial for a mechanistic understanding of attention because such rhythmic monitoring of locations or objects should gate the reallocation of attentional resources (Kastner & Buschman, 2017). To study the temporal fluctuation of attention, we used the attentional blink task, an RSVP two-target visual detection task. In these experiments, we discovered a perceptual phenomenon that we name *attentional rhythmic blink*: (1) second target (T2) detection performance in attentional blink fluctuated cyclically; (2) the presentation of distractors modulated the frequency of this behavioral oscillation. Specifically, detection of two targets in the presence of no distractors induced fluctuations of T2 detection accuracy at the theta frequency, accompanied by a prominent increase in the theta power in the parietal regions. Conversely, the presence of distractors induced fluctuations of T2 detection accuracy at the alpha frequency, accompanied by a prominent increase in the alpha power in the occipital regions. Our results provide compelling evidence that distractor suppression modulates the frequency of perceptual sampling.

In the conventional attentional blink, where T1-to-T2 lag is varied in steps of 100 ms (i.e., the lag is sampled at 10 Hz), second target detection was found to be deteriorated at T1-to-T2 lags of 300–500 ms. In contrast, we sampled T1-to-T2 lags on a much finer temporal scale (20 ms or 50 Hz, in Experiment 1; 40 ms or 25 Hz, in Experiment 2), which allowed us to capture the rapid fluctuations in behavioral performances. To the best of our knowledge, our study is the first to provide evidence of rhythmicity in the efficiency of detecting the second target in attentional blink performance, hence the novel term *attentional rhythmic blink*.

In our study, we have found that attentional rhythmic blink, in the absence of distractors, was driven at the theta rhythm (approximately 4 Hz in Experiment 1a). Theta rhythm in behavioral oscillation has been reported mainly in spatial attentional domain (e.g., Fiebelkorn et al., 2013), leading to the argument for a role of theta in spatial attentional exploration (sampling and shifting: Dugué & VanRullen, 2017; VanRullen, 2013). Quite differently, in our experiments, the two target stimuli were always presented at the same location but separated by a variable temporal interval, which required participants to allocate attention not spatially, but temporally. Thus, our results suggest a role for theta rhythm in attentional exploration more generally, both in the spatial and temporal domains (e.g., Zalta et al., 2020), supporting the idea that theta rhythm reflects general oscillatory mechanisms in multi-item attention (Huang et al., 2015; Re et al., 2019). Consistent with our behavioral data, the MEG experiment revealed an increase in theta power and a theta-phase dependence of the second target detection when no distractor was presented.

When distractors were presented temporally interspersed with the two targets, detection accuracy for the second target fluctuated at the alpha rhythm (approximately 8 Hz in Experiment 1b). Our MEG experiment showed that alpha power in the occipital region increased in the distractor condition, relative to the no-distractor condition (Experiment 2), suggesting a link to the attentional rhythmic blink at the alpha frequencies. This notion is consistent with a well-accepted framework proposing a role of alpha oscillations play a role in gating relevant information through functional inhibition (e.g., Jensen & Mazaheri, 2010, but see Foster & Awh, 2019). One earlier experimental support for this hypothesis is the increased occipital alpha power observed ipsilateral to the visually attended location (Kelly et al., 2006; Thut et al., 2006; Worden et al., 2000). Similar changes in alpha power have been reported in the domain of working memory (Bonnefond & Jensen, 2012; Jensen et al., 2002), leading to the idea that alpha oscillations reflect pulsed inhibition in sensory areas (Jensen et al., 2012; van Diepen et al., 2019). Adding further evidence in support of this general idea, our finding of increased occipital alpha power in the distractor condition suggests that alpha oscillations play a crucial role in suppressing task-irrelevant (distracting) inputs.

Given that rhythmic visual stimulation (e.g., 10 Hz visual flicker) modulates ongoing rhythmic neural activity and, accordingly, subsequent visual performance (de Graaf et al., 2013; Helfrich et al., 2017; Helfrich et al., 2019; Spaak et al., 2014), the alpha power increase in the distractor condition might originate from sensory entrainment by RSVP streams, rather than active inhibition of distractors. Although we presented distractors with jittered intervals to reduce the entrainment effect (see Fig. S5 for presentation frequency distributions), jittered presentation might also entrain alpha (Mathewson et al., 2012; Zauner et al., 2012, but see Cravo et al., 2013). However, we believe that entrainment alone cannot account for our results. If such repetitive visual stimulation was the sole cause to induce alpha sampling in our study, then the same alpha rhythm and power should be observed, independent of T1 detection. However, analysis of T2|T1-*error* trials in Experiment 1b (i.e., conditional analysis of T2 detection in trials of T1 errors) showed not alpha, but theta peak in their performance (Figure S2), even though visual inputs in T2|T1-correct and T2|T1-error trials were the same (Figure 1). We thus argue that the observed alpha sampling in our experiments is derived from distractor suppression, not solely alpha entrainment due to visual stimulation. Future work may be able to further rule out the potential confound by presenting stimuli at a higher frequency than alpha (e.g., Martin et al., 2011) or equalizing visual displays by employing weak and strong distractors in RSVP streams. Furthermore, while we here interpret that alpha power increase and rhythmic sampling is associated with suppression of distracting items in RSVP streams, one alternative interpretation might be that these are rather associated with successful maintenance of T1 in working memory (e.g., Bonnefond & Jensen, 2012). Future study should explore whether the increase in alpha power is driven by ignoring distracting visual inputs or maintenance in working memory, or both.

Based on the current results, we propose a unified model of distinct contributions of theta and alpha rhythms: the power balance of theta and alpha oscillations determine the frequency of behavioral oscillations. The model is an extension of rhythmic attention model in Dugué and VanRullen (2017), claiming that the natural alpha rhythm at the early visual cortex determines the perceptual sensitivity while the rhythmic feedback (∼5 Hz) from higher-order region to sensory areas enables theta-rhythmic attentional exploration. At the onset of visual events, feed-forward signals are sent from the occipital area to higher-order regions, which causes periodic feedback to the sensory area to produce rhythmic attentional exploration. Supporting this view, attentional theta rhythm has been reported to originate from the front-parietal attentional network (Helfrich et al., 2017, 2018; Fiebelkorn et al., 2019). In the no-distractor condition of the current study, presentation of the first target (T1) would produce feed-forward activation and rhythmic feedback, resulting in more dominant parietal theta over occipital alpha as reflected in pre-T2 power change (Fig 4a). Such a theta-dominant neural state could produce theta sampling in visual perception (Fig 5a left) as observed in attentional rhythmic blink at the theta (Experiment 1a) as well as the T2 detection dependent on the parietal theta phase (Experiment 2). In the distractor condition, on the contrary, observers should suppress task-irrelevant information to select the target, which would require an alpha power increase for inhibition. In fact, we found alpha power increase in the occipital area, accompanied by the attentional rhythmic blink at alpha (Experiment 1b) as well as the T2 detection dependent on the occipital alpha phase (Experiment 2).

**Fig. 5:**
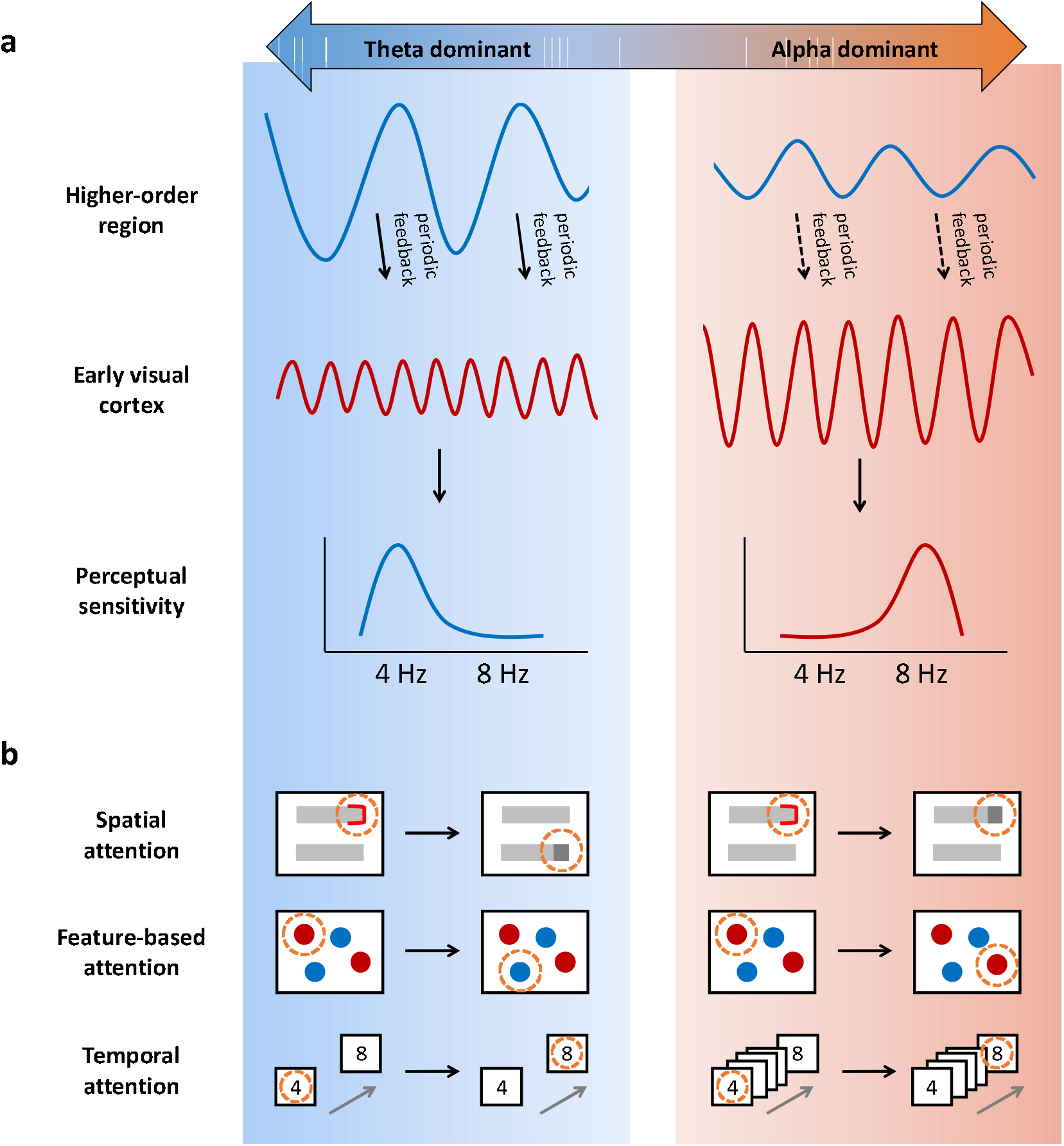
Tentative model of theta- and alpha-rhythmic sampling. (extending the model by Dugué and VanRullen (2017)). **a** On the left, periodic feedback from higher-order region to sensory area results in theta-cyclic modulation of visual sensitivity. On the right, the alpha power at the early visual cortex may become dominant over theta (due to distractor suppression, for example), and then the perceptual sensitivity fluctuates in the alpha instead of theta-band. Thus, the dominance of neural oscillations determines the perceptual threshold. **b** Realization of rhythmic multi-item sampling in spatial attention (e.g., Fiebelkorn et al., 2013), feature-based attention (e.g., Re et al., 2019), and temporal attention, as reported in the current study.

The neural balance between theta and alpha power might be able to explain not only the current results but also the reported frequency changes in behavioral oscillations such as spatial attention (Fiebelkorn et al., 2013) or feature-based attention task (Re et al., 2019). Namely, in those studies, behavioral performance fluctuated at theta when multiple locations or features should be explored (left panels of Figure 5b), while it oscillated at alpha when neural representations should be protected from interference of other locations or features (right panels of Figure 5b). We here argue that the frequency of behavioral rhythms depends on the needs of the change in brain representations or its protection from interference. Consistent with this idea, updating neural representations from current to alternative dimensions, that is top-down switching process, has been associated with theta power increase (Sauseng et al., 2006). Critically, at the preparation stage for task switching, increased theta and decreased alpha power over occipital areas were observed (Gladwin & de Jong, 2005). This theta power increase accompanied by cognitive engagement of updating representation for shifting attention should be the base for reported theta sampling for detecting the stimuli that are different from cued location or cued color. In contrast, protection of neural representations from interference has been associated with alpha power increase (e.g., Jensen & Mazaheri, 2010). In fact, modulatory effects of alpha power are not limited to spatial locations (e.g., Thut et al., 2006), but have also been reported for feature-based attention (e.g., Snyder & Foxe, 2010). This alpha power increase should be a key for reported alpha sampling for detecting the stimuli that at the cued location or of the cued color. In sum, theta oscillations for exploration and alpha oscillations for maintenance have complementary roles for information processing, and their balance determines the behavioral rhythms in visual perception (Gratton, 2018).

To summarize, we have demonstrated that suppression of visual distractors modulates rhythmicity in visual perception. When two targets but no distractors were presented, we observed behavioral rhythms at the theta frequencies, accompanied by an increase in theta power over the parietal area, and detection accuracy depended on its phase. When two targets were embedded in a temporal sequence of distractors, detection accuracy of the second target was rhythmically fluctuated at alpha frequencies, accompanied by an increase in alpha power over the occipital area, and detection accuracy depended on its phase. We propose that the frequency of perceptual rhythm is determined by the balance of multiple neural rhythms associated with task demands. We believe that our model provides a new conceptual framework to guide further investigation into how perceptual rhythm is determined in a complex and noisy natural environment.

## Methods

All the experiments were approved by the local ethics committee at the National Institute of Information and Communications Technology.

## Behavioral Experiment

### Participants

Thirty-one individuals (10 females), age 20–26 years (mean = 22.5, *SD* = 1.5) participated in the no-distractor experiment (Experiment 1a), and 30 individuals (15 females), age 20–25 years (mean = 22.5, *SD* = 1.5) participated in the distractor experiment (Experiment 1b). In the latter, two subjects were removed because their performances on T1 detection were below the chance level, reducing the number of subjects to *n* = 28. All participants were right-handed and reported normal or corrected-to-normal visual acuity. Participants gave informed consent before participating in the experiments and received monetary compensation (3,000 Japanese Yen, the equivalent of approximately 28 US dollars).

### Apparatus and stimuli

The stimuli were presented on a 27-inch display monitor (ASUS ROG PG278Q) using MATLAB (Mathworks Inc. Natick, MA) and Psychtoolbox (Braianard, 1997; Pelli, 1997). The screen had a resolution of 1920 × 1080 pixels, at a refresh rate of 100 Hz. The viewing distance was fixed at approximately 70 cm with a chin rest. Subjects’ responses were collected using a numerical keypad.

In both experiments, all stimuli were presented on a gray background (RGB: 128, 128, 128). The first target (T1) and the second target (T2) were in Courier New font, spanned a 1° × 1° visual area, and were presented for 30 ms. Both were masked with an image of random black and white line-fragments presented for 30 ms immediately after the target. T1 and T2 were randomly chosen from the digits 2,…,9 without duplication.

### Design and procedure

#### No-distractor condition (Experiment 1a)

Immediately after participants pressed the enter key, a black fixation cross (0.5° × 0.5°) was presented for 500, 600, or 700 ms. After that, in each trial, two targets were presented, but no distractors. Participants had to detect the two successive targets (minimalist attentional blink or skeletal attentional blink task: e.g., Duncan et al., 1994; McLaughlin et al., 2001). Critically, the T1-to-T2 lag was set between 100 ms and 1000 ms, randomly sampling this interval discretized in 20 ms steps (corresponding to 50 Hz sampling rate), providing 46 distinct realized intervals. At the end of each trial, participants were asked to report the identities of T1 and T2 with a key press. They were instructed to respond as accurately as possible with no time pressure, and feedback of their key presses (correct/incorrect) were provided at the end of each trial for 300 ms. Inter-trial intervals were set at 800 ms. To obtain a more noticeable effect of cue resetting, the number of trials for SOA of 100 ms was 10 times of that for other SOAs (Fiebelkorn et al., 2011; Song et al., 2014). The main experiment consisted of four blocks of 220 trials, for a total of 880 trials (i.e., trials of SOA of 100 ms contained 160 trials and others included 16 trials). The experiment was preceded by 20 practice trials.

#### Distractor condition (Experiment 1b)

The design and procedure were the same as in Experiment 1a, with the following exception. After the fixation period, the RSVP streams were presented (30 ms/item) with jittered inter-item intervals (randomly selected from the 70 to150 ms interval discretized in 10-ms steps). The purpose of this manipulation of jittered inter-item intervals was not to present items rhythmically at a specific rate to avoid neural entrainment.

### Behavioral data analysis

#### Classic attentional blink

Smoothing, using ten-point moving average, was applied to the raw accuracy time course for the T2|T1-correct trials. We compared the difference of the smoothed accuracy at the lags of 300 ms and 800 ms. A significant increase from 300 ms to 800 ms was used to verify that the classic attentional blink phenomenon was observed.

#### Accuracy fluctuations

The temporal profile of the accuracy was calculated as a function of the T1-to-T2 lag. Each participant’s detection accuracy was detrended using a second order polynomial. We then performed discrete Fourier transform to convert the temporal detection accuracy function into the frequency domain (using zero padding with a Hanning window). Amplitude spectra were derived by taking the absolute value of the complex FFT output. We performed a randomization procedure by shuffling the time points of the accuracy time series to assess statistical significance of peaks in the spectra amplitude. Each randomization was applied for each participant separately. After each randomization of accuracy time series, the FFT was performed exactly as on the unshuffled data. This was repeated 1000 times, producing a distribution of spectral power for each frequency point from which we obtained the significance threshold (*p* < 0.05; uncorrected). Multiple-comparison correction was performed by setting the maximum across all frequency bins as the threshold (single threshold test; Nichols & Homes, 2002; Huang et al., 2015; Song et al., 2014).

## MEG Experiment

### Participants

In Experiment 2, 30 undergraduate and graduate students participated in the experiment as volunteers or for monetary compensation (8,000 Japanese Yen, or approximately 74 US dollars). Moreover, four of them participated in Experiment 1a and five of them participated in Experiment 1b. One was removed because of chance level performances on T1 detection in the no-distractor condition, and two were removed because of their extremely high-performance levels in the T2|T1-correct trials in no-distractor condition (>96%; more than 16 of 23 data points showed 100% accuracy). In total, 27 participants (8 females), age 21–29 years (mean = 22.9, *SD* = 1.9), were included in the analyses. All were right-handed and reported normal or corrected-to-normal visual acuity.

### Apparatus and stimuli

Visual stimuli were projected onto a screen in the MEG scanner, in front of the participant, with a Digital Light Processing projector (Panasonic model PT-DZ630) operating at 60 Hz refresh rate. The viewing distance was approximately 57 cm. Stimulus onset was recorded using photodiodes located at the top-right corner of the screen.

### Design and procedure

Each target was presented for 33.3 ms, followed by mask stimuli for 33.3 ms. T1-to-T2 lag was set between 116.7 ms to 850 ms, in steps of 33.3 ms (23 data points, corresponding to 25 Hz sampling rate). Participants were asked to report the two numbers from four choices as accurately as possible with no time pressure (i.e., chance level is 25%). They were instructed to respond as accurately as possible with no time pressure, and feedbacks of their key presses were provided at the end of each trial for 300 ms.

The two types of trials (no-distractor and distractor trials) were blocked and presented in a counterbalanced order across participants (half of them completed no-distractor trials first, then distractor trials). The main experiment consisted of six blocks of 108 trials, with the opportunity for a brief rest every 36 trials, for a total of 648 trials (i.e., trials of SOA of 116.7 ms contained 60 trials and others included 12 trials). The experiment was preceded by 20 practice trials for both the no-distractor and distractor conditions.

### MEG scanner and data acquisition

MEG data were collected with Elekta Neuromag 306 channel system (Elekta Neuromag? 360TM, MEGIN/Elekta Oy, Finland; 102 magnetometers, 204 planar gradiometers) with a sampling frequency of 1 kHz (0.03–330 Hz bandpass filter). Prior to the recording, four head position indicator (HPI) coils were attached to the participant’s forehead. A digitizer device was then used to record the locations of fiducial points (right and left pre-auricular points and nasion), the HPI coils, and their head surface points. A 2-min empty-room recording was obtained before the experiment. This data was used to estimate sensor and environmental noise statistics for MEG source modeling.

### Behavioral data analysis

#### Classic attentional blink

Smoothing, using ten-point moving average, was applied to the raw accuracy time course for T2|T1-correct trials. A significant increase accuracy from the lag of 316.7 ms to 716.7 ms was used to verify that the classic attentional blink phenomenon was observed.

#### Accuracy fluctuations

The same analysis as in Experiment 1 was performed.

### MEG data analysis

#### Preprocessing and HPI correction

Raw MEG data was first preprocessed using the oversampled temporal projection method, which suppresses uncorrelated sensor noise and artifacts (OTP: Sig3 Denoise-OTP™, Sigma3 SP LLC, US; Larson & Taulu, 2018). The MEG data was then preprocessed using MaxFilter software (version 2.2.12, MEGIN/Elekta Oy, Finland). Bad channels were automatically identified with Xscan (version 3.0.15, MEGIN/Elekta Oy, Finland) and were removed from the subsequent calculations for environmental noise. We then performed spatiotemporal signal space separation (tSSS: Maxfilter-2.2™, MEGIN/Elekta Oy, Finland; Taulu & Simola, 2006) using the remaining channels with a sliding window of 10 s and a correlation limit of 0.980, with the origin of the head set to (0, 0, 40 mm). To correct for head movements between sessions, MEG signals were transformed to the participant’s head-centered coordinates at the beginning of the first session using the –trans option of MaxFilter.

#### Artifact removal

All further data analysis was done using MATLAB (Mathworks Inc. Natick, MA) and the Brainstorm open-source software (Tadel et al., 2011). An offline digital notch filter with the center frequency of 60, 120, and 180 Hz was applied to remove the power line interference. Using Independent Component Analysis, eye-related and heartbeat artifacts were identified and removed.

#### Source space modeling and HPI-MRI alignment

Head scans, using high-resolution T1-weighted anatomical MRIs (3.0 T Trio, Siemens), were collected from each participant. Cortical surfaces were extracted from each structural MRI using CAT12 (computational anatomy toolbox: Dahnke et al., 2013) for SPM12 (Statistical Parametric Mapping software, http://www.fil.ion.ucl.ac.uk/spm/). A 15,000-vertex cortical surface source model was used for head modeling. Using the locations of the HPI coils measured on the scalp and the fiducial points, the MEG sensor space was co-registered to the participant’s own MRI space.

We obtained an MEG forward model using the overlapping-spheres approach (Huang et al., 1999). Depth-weighted minimum norm estimates (DMNE) were then performed separately for each subject using Brainstorm software. The weighted-minimum norm operator included an estimate of the variance of noise at each MEG sensor obtained from the empty-room recording. Trial level source activity was computed using dSPM (dynamic statistical parametric mapping). Cortical activations were analyzed in each participant using anatomically defined regions of interest (ROI). We selected these regions from the Desikan–Killiany atlas (Desikan et al., 2006). Based on observations in previous reports (Dugué & VanRullen, 2017; Helfrich et al., 2017, 2018; Fiebelkorn et al., 2019), we focused our analysis on superior parietal and lateral occipital ROIs in both the left and right hemispheres.

#### Time-frequency analysis of pre-T2 onset

MEG source signals were transformed to time-frequency representations with Morlet wavelet transform (1000 ms pre-T2 to 1000 ms post-T2 time; center frequency = 1 Hz, FWHM = 3 s). To assess and isolate the contribution of distractors preceding the T2 presentation, analysis was restricted to an 800 ms time window (400 ms pre-T2 to 400 ms post-T2). Data thus segmented were baseline corrected (as the percentage change from baseline defined in the −500 to −400 ms interval), which resulted in an estimate of oscillatory power at each time sample and at each frequency between 3 and 40 Hz. We performed 1000 permutation tests to compare the time-frequency patterns between distractor and no-distractor conditions for each ROI.

#### Phase analysis

We explored the effect of pre-stimulus phase on T2 detection accuracy by focusing on identified frequency bands. Band-pass filtering for single-trial event-related fields in theta and alpha bands was applied. The oscillatory phase was calculated using the Hilbert transform of the filtered signals separately in trials grouped by accuracy (correct vs. incorrect) and condition type (distractor vs. no-distractor). The reference time-point for each frequency band was defined by the time of observed minimum *p*-values (maximum significance) for the power (Dugué et al., 2011). If successive time points were detected, the median of the time points was used for time-point of interest. Phase distributions for each frequency band for each condition type was compared, correct versus incorrect trials, using the Watson-Williams circular test (CircStat toolbox; Berens, 2009).

## Code availability

All custom code used for this manuscript is available upon request from the Lead Contact: Tomoya Kawashima.

## Data availability

All data contained in this manuscript are available upon request from the Lead Contact: Tomoya Kawashima.

## Author Contributions

T.K. performed experiments and analyzed data. M.J.H, K.A., and T.K. designed experiments, defined, and validated data analysis methods, and wrote the paper.

## Competing interests

The authors declare no competing interests.

## Corresponding author

Correspondence to Tomoya Kawashima.

## Acknowledgements

We thank Asuka Otsuka, Takeshi Nogai and Hironori Nishimoto for expert help in MEG recordings. We also thank Yasuki Noguchi, Nobuhiro Hagura, Hiroshi Ban, and Ryohei Nakayama for providing critical discussions and comments on an early draft. The research was supported by JSPS Grant-in-Aid for Scientific Research on Innovative Areas, “Chronogenesis: how the mind generates time” (No. 8002) to K.A.; JSPS Grant-in-Aid for Research Activity Start-up (JP18H05813, JP19K21005) and JSPS Grant-in-Aid for Early-Career Scientists (JP20K14274) to T.K.; JSPS Grant-in-Aid for Scientific Research JP18H01101, Grant-in-Aid for Scientific Research on Innovative Areas JP19H05313 and JP21H00315 and JST PRESTO JPMJPR19J8 to M.J.H.

## Notes

### Competing Interest Statement

The authors have declared no competing interest.

