## Supplementary File for "Attentional rhythmic blink: Theta/Alpha balance in neural oscillations determines the rhythmicity in visual sampling"

**Supplemental Information**

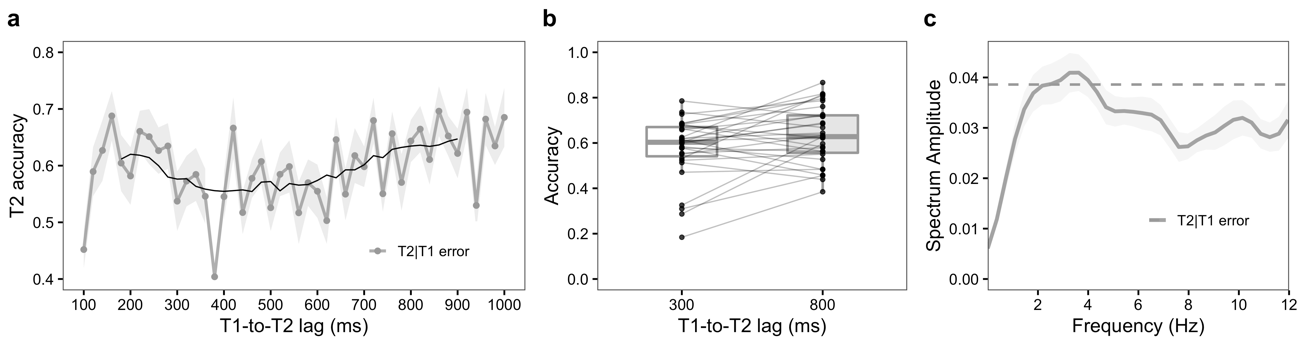

**Fig. S1: Attentional blink and its rhythms with no distractor (Experiment 1a) for T2|T1-error trials (*n* = 31).** **a** Grand average accuracy time course (mean ± SEM) as a function of T1-to-T2 lag (sampled in 20 ms steps) for T2|T1-error trials. Smoothing with a ten-point moving average (solid line) was used to approximate the classic attentional blink phenomena. **b** The mean T2 detection accuracy obtained after the smoothing for each subject (dots) inT2|T1-error trials. Overall, accuracy was higher at lag 800 ms (63%) than at lag 300 ms (58%) (*t* (27) = 2.58, *p* = 0.015, *d* = 0.46), consistent with the classical attentional blink effect. **c** Grand average spectrum (mean ± SEM) for detrended accuracy time courses, as a function of frequency in the 0 to 12 Hz band for the T2|T1-error condition. The dashed line indicates the statistical significance threshold (*p* < 0.05) after permutation test and corrections for multiple comparisons.

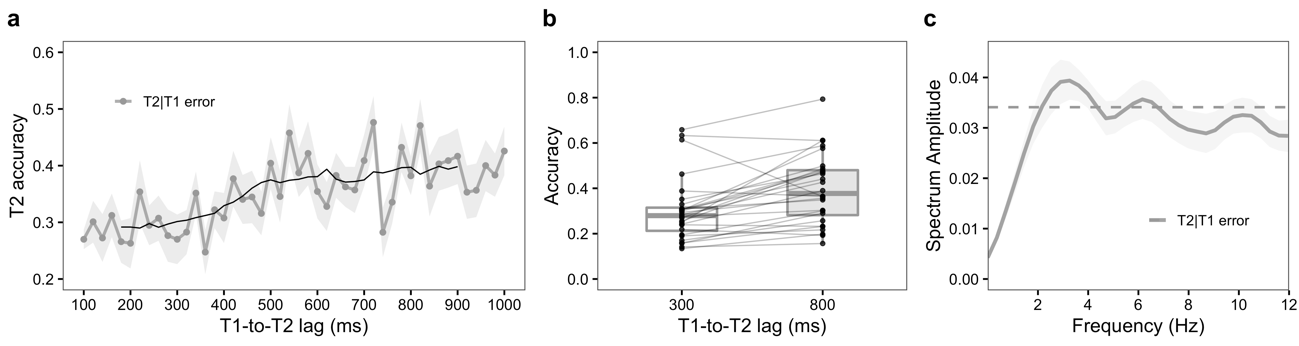

**Fig. S2:** **Attentional blink and its rhythms with distractors (Experiment 1b) for T2|T1-error trials (*n* = 28 subjects).** **a** Grand average accuracy time course, constructed as in Fig. S1. **b** The mean T2 detection accuracy, constructed as in Fig. S1. Accuracy was higher at lag 800 ms (40%) than at lag 300 ms (30%) (*t* (27) = 4.42, *p* <0.001, *d* = 0.84), consistent with the classical attentional blink effect. **c** Grand average spectrum, constructed as in Fig. S1.

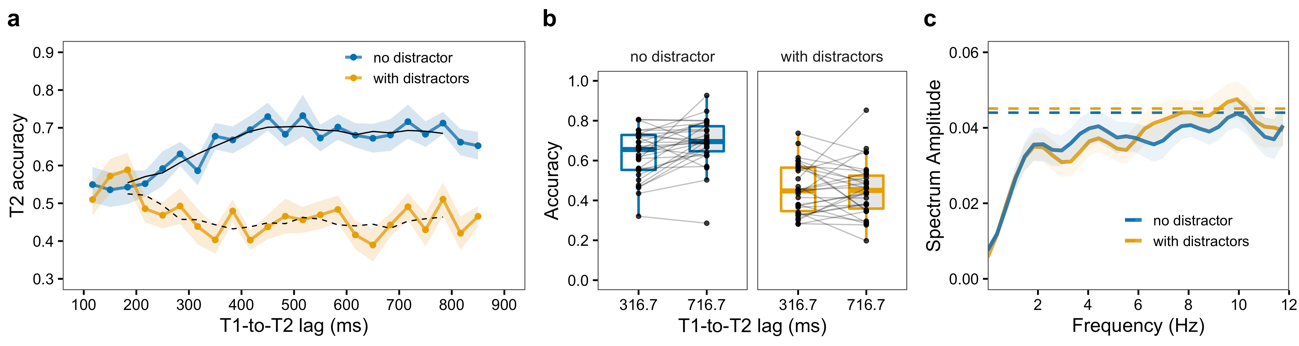

**Fig. S3:** **Attentional blink and its rhythms in MEG experiments (Experiment 2) for T2|T1-correct trials (*n* = 27).** **a** Grand average accuracy time course (mean ± SEM) as a function of T1-to-T2 lag (sampling the 116.7 ms to 850 ms interval in steps of 33.3 ms), for distractor (yellow) and no-distractor (blue) conditions. Smoothing with a five-point moving average (solid for distractor and dashed for no-distractor conditions, respectively) was used to approximate the classic attentional blink phenomena. **b** The mean T2 detection accuracy obtained after 5-point moving average for each subject and trial type. Each dot represents the mean accuracy for one subject. For two lags (316.7 ms or 716.7 ms) and 2 conditions (with or without distractor), we performed a 2 × 2 repeated measure analysis of variance (ANOVA) after the 5-point smoothing of the raw accuracy time courses (Figure 4). Main effect of condition type and lag were both significant (*F* (1, 26) = 71.91, *p* < 0.001, $\eta_{p}^{2}$ = 0.73; and *F* (1, 26) = 9.13, *p* = 0.006, $\eta_{p}^{2}$ = 0.26, respectively), showing that the overall accuracy was higher for no-distractor condition. Interaction between lag and trial type was significant (*F* (1, 26) = 4.70, *p* = 0.040, $\eta_{p}^{2}$ = 0.15), which reflected a lag effect (i.e., the classical attentional blink phenomenon) for the distractor condition (*F* (1, 26) = 13.62, *p* = 0.001, $\eta_{p}^{2}$ = 0.34) but not for no-distractor condition (*F* (1, 26) = 1.02, *p* = 0.323, $\eta_{p}^{2}$ = 0.04). **c** Grand average spectrum (mean ± SEM), constructed as in Fig. S1. The dashed lines (also constructed as in Fig. S1) indicate the corresponding statistical significance threshold (*p* < 0.05). When distractors were presented, T2 accuracy fluctuated at alpha frequencies (6.3–7.0 Hz: *p* < 0.05, corrected for multiple comparison), but showed no specific dominant frequency in its fluctuations in the absence of distractors. The low number of trials used here (as compared to Experiment 1) may explain the lack of significance. These results support the idea that the presence of distractors modulates the rhythm of perceptual efficacy.

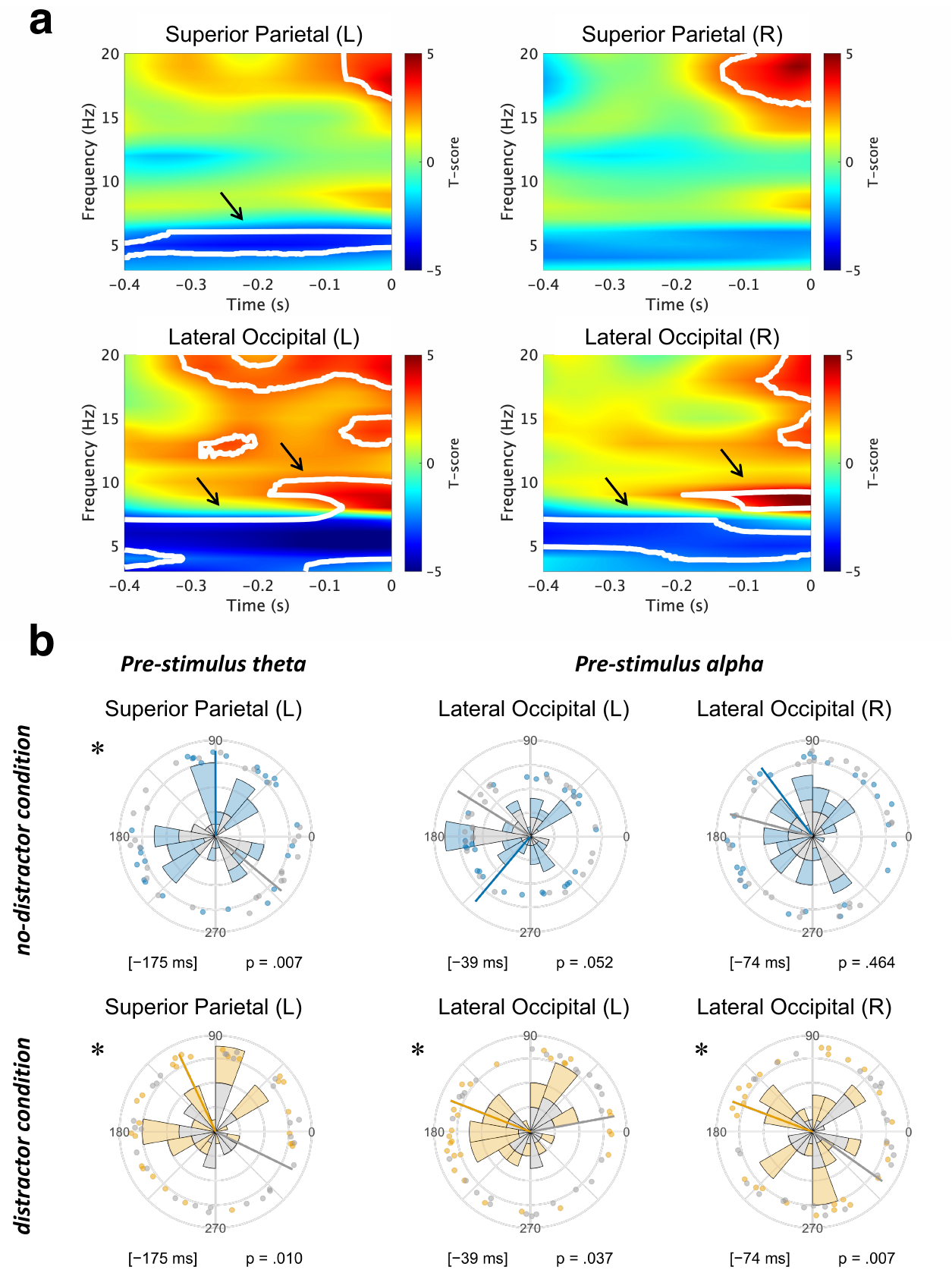

**Fig. S4: Pre-stimulus oscillatory power and phase differences in Experiment 2**. No-distractor and distractor conditions were presented block by block. Data from the left hemisphere are also presented in Fig 4 and the main text. **a** Normalized time-frequency difference spectra (distractor minus no-distractor condition) simultaneously recorded in left (L) and right (R) superior parietal and lateral occipital areas and averaged across subjects (*n* = 27). Time is relative to T2 presentation. White contours denote significant cluster-corrected effects (*p* < 0.05, FDR corrected). Note the distractor effects at pre-stimulus times, suppression (blue) in the theta (4–7 Hz) and enhancement (red) in the alpha (8­–10 Hz) frequency bands (arrows). **b** Phase analysis of the pre-T2 MEG signals recorded in the theta-band (4–7 Hz) and alpha-band (8–10 Hz) for no-distractor (top) and distractor (bottom) conditions. Data in color are from trials of correct T2-detection; those in gray, for incorrect T-2 detection. Each dot represents the mean phase angle per participant, wedges represent the circular phase distribution in the subjects (thick lines represent the mean phase angle across subjects). Distractor vs. no-distractor phase differences were analyzed using a circular Watson-Williams test. The asterisks indicate significant phase difference (*p* < 0.05).

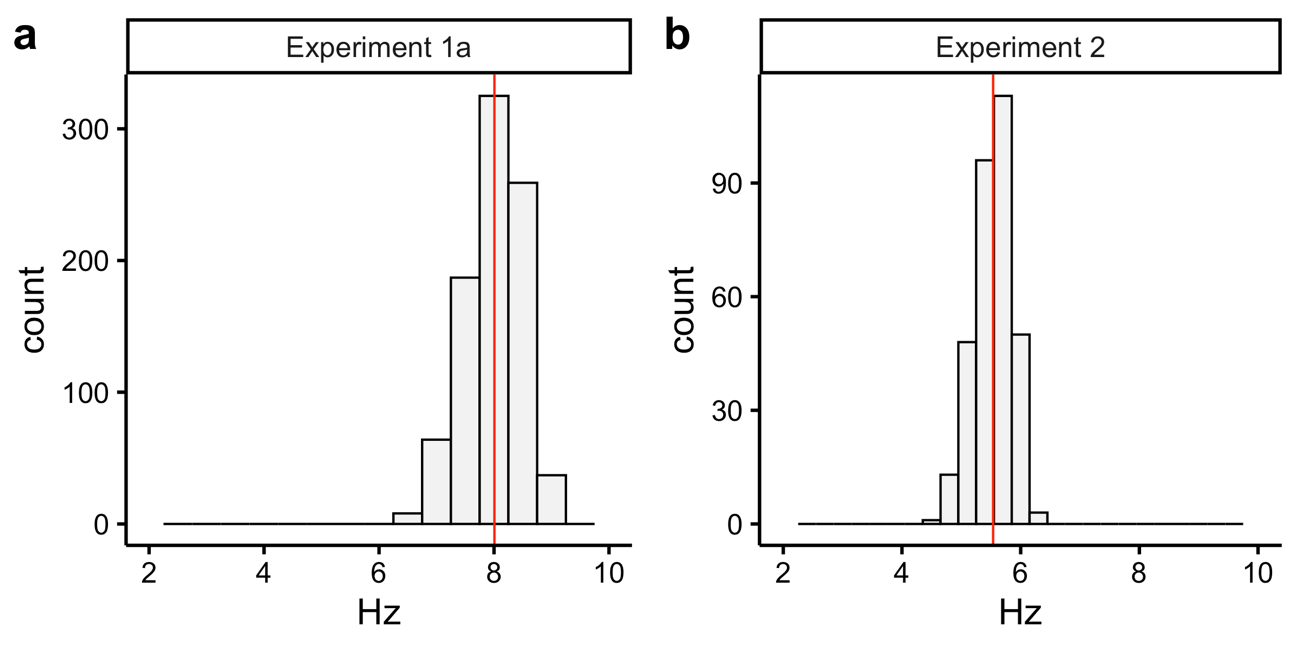

**Fig. S5: Distribution of the presentation frequency.** We calculated the average presentation frequency of RSVP streams per trial. Red lines indicate the mean. **a** Experiment 1a, no-distractors condition, mean (±SD) at 8.0 ± 0.5 Hz (*n* = 880 trials). **b** Experiment 2, distractor condition, 5.5 ± 0.31 Hz (*n* = 648 trials).
